# Integrated Patient-Derived Xenograft and Patient-Derived Cell Models Reveal Therapeutic Vulnerabilities Beyond Standard-of-Care Therapy in Endometrial Cancer

**DOI:** 10.64898/2026.07.24.740568

**Authors:** Tianyue Li, Fang Huang, Xiaohao Huang, Ella Pate, Riley Rosenmeyer, Mya Messenger, Kaylee McSweeney, Samuel Robinson, Adam Deters, Lauren Buchanan, Maggie Meehan, Nandini Patel, Allison Diekema, Yiqin Xiong, Xiaofang Zhang, Xiangbing Meng, Shujie Yang

## Abstract

Endometrial cancer (EC), the most common gynecologic malignancy in the USA, has seen limited improvement in patient outcomes over recent decades, underscoring the need for relevant preclinical models. To address EC heterogeneity, we established an integrated platform of patient-derived xenografts (PDXs) and matched patient-derived primary cancer cells (PDCs) for disease modeling and systematic drug sensitivity testing.

Fresh tumor specimens (n=103) were collected from EC patients to generate PDXs in immunodeficient mice and corresponding PDCs. Fifty-three PDX models were successfully established (52% engraftment rate), with higher success observed in high-grade, recurrent, metastatic tumors (70%), compared with their low-grade counterparts (56%). Histopathologic and immunohistochemical analyses confirmed that PDX tumors faithfully preserved morphology, hormone receptor status, and intertumoral heterogeneity across multiple passages.

Using 13 PDC models, we performed an unbiased screening of 179 FDA-approved oncology drugs, revealing marked intertumoral variability in drug response. Almost all PDC models exhibited limited sensitivity to NCCN-recommended therapies, highlighting the need for alternative treatment strategies. In contrast, multiple FDA-approved agents including epigenetic modulators, dual PI3-kinase/HDAC inhibitors, topoisomerase II inhibitors, and proteasome inhibitors demonstrated potent antitumor activity. Importantly, a low-dose combination of the DNA methyltransferase inhibitor 5-azacytidine and the histone deacetylase inhibitor romidepsin significantly suppressed tumor growth across six independent PDX models.

Together, these findings establish a comprehensive PDX and PDC platform as a robust translational resource. By capturing the histopathologic and molecular diversity of EC and identifying clinically actionable therapeutic advantages, including an epigenetic combination regimen, this study offers a translational resource for preclinical drug evaluation.

## Introduction

Endometrial cancer (EC) is the most common gynecological malignancy in the United States and is the only cancer type with declining survival rates over the past 40 years [1]. In 2026, the NIH estimates 68,270 new cases and 14,450 deaths from EC, reflecting a growing public health burden driven by rising incidence and mortality rates [2]. Despite advances in surgical techniques and systemic therapies, the 5-year survival rate for EC has decreased from approximately 87% to 81% over the past 40 years [3], underscoring the urgent need for improved therapeutic strategies and biologically informed treatment paradigms.

EC is a highly heterogeneous disease characterized by diverse histologic subtypes and molecular landscapes that profoundly influence therapeutic response and clinical outcomes. This heterogeneity complicates treatment selection and contributes to variable responses to standard-of-care therapies, including surgery, hormonal therapy, chemotherapy, radiotherapy, immunotherapy, and targeted therapies [4]. Robust preclinical models are essential to advance scientific understanding of EC biology and to bridge the gap between molecular characterization and therapeutic decision-making [5]. There is still a limited number of viable models that faithfully represent EC patients, driving us to work to fill this gap [6].

Patient-derived models, such as patient-derived xenografts (PDX) and patient-derived primary cancer cell lines (PDC), offer a powerful alternative by closely recapitulating human tumors [7, 8]. These models preserve key histopathological features, genomic alterations, and molecular signaling networks of the original patient tumors, enabling the study of tumor evolution, heterogeneity, and drug response in a clinically relevant context. By maintaining inter- and intra-tumoral diversity, PDX and PDC models provide a critical platform for understanding how distinct tumor subpopulations contribute to disease progression and therapeutic resistance.

In this study, we establish and characterize a comprehensive platform of patient-derived EC models, including matched PDX and PDC systems, designed to recapitulate the morphological, molecular, and biological features of primary tumors. By preserving tumor heterogeneity and patient-specific tumor behavior, these models enable in-depth investigation of EC biology and treatment response.

Furthermore, PDX and PDC models were utilized to evaluate the efficacy of anticancer drugs, identifying key factors influencing therapeutic responses that inform treatment strategies. Unlike prior EC-PDX studies focused on molecular profiling or single-agent testing, our study integrates matched patient-derived cells with unbiased FDA-approved oncology drug screening and *in vivo* validation [9]. This combined strategy establishes a functional precision medicine platform for EC and highlights the importance of integrated PDX and PDC models for functional drug testing and therapeutic evaluation in EC [10, 11].

## Methods

### Mice

Nonobese diabetic/severe combined immunodeficient (NOD/SCID) gamma (NSG; NOD.Cg-Prkdcscid Il2rgtm1Wjl/SzJ, Cat. #005557) and green fluorescent protein-expressing (GFP)-NSG (NOD.Cg-Prkdcscid Il2rgtm1Wjl Tg (CAG-EGFP)1Osb/SzJ, Cat. #021937) mice were purchased from Jackson

Laboratory and bred at the University of Iowa animal facility. Mice were kept in a controlled environment (22°C, 50% relative humidity, 12-hour light-dark cycle) under specific pathogen-free (SPF) conditions. All experiments were approved by the Institutional Animal Care and Use Committee (IACUC) and followed the Guide for the Care and Use of Laboratory Animals, the Animal Welfare Act, and the Public Health Service (PHS) guidelines (PHS Assurance No. D16-00009 [A3021-01]).

Only female mice were used, as endometrial cancer occurs in biological females.

### Patients

Between December 2022 and September 2025, a total of 103 endometrial cancer cases were collected. The study was approved by the Institutional Review Board at the University of Iowa (IRB #201708847), and informed consent was obtained from all patients. All patient data was collected, managed, and analyzed in accordance with ethical guidelines.

### Hematoxylin and eosin (H&E), Immunohistochemistry (IHC) staining, and Endometrial Cancer Mutation Profile (ECMP)

Following receipt of tumor tissue, a 1-2 mm cross section was obtained under sterile conditions and fixed in 10% neutral-buffered formalin. Subsequently, formalin-fixed, paraffin-embedded (FFPE) tissue blocks were prepared, and H&E, IHC, and immunofluorescence analyses were performed by the Tissue Procurement Core in the Department of Pathology at the University of Iowa. The same analyses were conducted on PDX tumor tissue over multiple generations. ECMP testing was conducted and interpreted by the Molecular Pathology Division, Department of Pathology, University of Iowa.

### PDX Models

Matched normal and tumor tissue was provided by the University of Iowa Hospitals & Clinics (UIHC) following surgical removal from patient endometrium. Tumor fragments, (∼0.1 x 0.1 x 0.2 cm) were subcutaneously implanted bilaterally into the flanks of mice under sterile conditions. This was done for all EC subtypes except for hyperplasia. Tumor size was measured regularly using a caliper. Per IACUC guidelines, mice were euthanized, and tumors were harvested prior to reaching 20mm in length. Tumor volumes were calculated as length × width² × 0.5. Tumor samples were processed for dissociation, followed by two-dimensional (2D) and three-dimensional (3D) culture, and were snap-frozen for downstream analyses.

### PDC Models

All cell culture experiments were performed under sterile conditions. Tumors were excised, minced into small fragments, and digested in pre-warmed DMEM/F12 (Gibco, Cat. #11320033) containing 0.5 mg/mL collagenase A (Sigma-Aldrich, Cat. # 10103578001) and 20 μg/mL of DNase I (Sigma-Aldrich, Cat. #10104159001) at 37°C while shaking for 30 minutes. The resulting suspension was filtered through a 70 μm nylon mesh strainer (Fisherbrand) and washed twice with PBS. The filtrate was centrifuged at 1,200 rpm for 5 minutes, and the cell pellet was retained. The remaining tissue was further dissociated in pre-warmed 0.25% trypsin-EDTA (Gibco, Cat. # 2520056) and Accutase (Sigma-Aldrich, Cat. #A6964) (1:1) at 37°C while shaking for 10 minutes. The digestion was stopped by adding 10 mL of complete medium (DMEM/F12 supplemented with 10% FBS (R&D Systems, Cat. #S11150)), and the suspension was centrifuged at 1,200 rpm for 5 minutes before plating in TC60 dishes in 6 mL of medium (DMEM or DMEM/F12 supplemented with 1% penicillin-streptomycin (Gibco, Cat. #151400122) and 0.5% gentamicin (Research Products International (RPI), Cat. #G38000-5.0) and 5% or 10% FBS). Cultures were maintained at 37°C in a 5% CO_2_ incubator and routinely checked for mycoplasma contamination and confluency.

### 3D Matrigel spheroid

A 3% Matrigel solution (Corning, Cat. #354234, 8.6 mg/mL) was prepared and added to 6-well plates (VWR, Cat. #10028-028) followed by incubation at 37℃ to allow solidification. A total of 8 × 10^6^ cells were seeded per well to reach ∼80% confluency after 8 days and harvested. For collection, the growth medium was aspirated, and the wells were rinsed twice with ice-cold PBS. Matrigel was dissolved by adding ice-cold PBS-EDTA, supplemented with 5 mM EDTA and 1x proteinase and phosphatase inhibitor cocktail (Fisher Scientific, Cat. #78443) and shaken at 4°C for 30 minutes. The suspension was centrifuged at 3,000 rpm for 5 minutes, the supernatant was partially removed, and the colonies were resuspended in the remaining volume. For immunofluorescence analysis, coverslips were coated with Poly-D-Lysine (Sigma-Aldrich, Cat. #P6407) prior to plating the colony suspension. Cells were fixed with 4% PFA, permeabilized with 0.2% Triton X-100 (Fisher Scientific, Cat. # AAA16046AE) and blocked with 3% BSA/PBS (RPI, Cat. #C802G45) before labeling with primary antibodies against PR, followed by Alexa 488-conjugated goat anti-rabbit secondary antibody (Supple. Table 1).

### FDA oncology drug screening

High-throughput drug screening was performed on 13 PDC models. 96-well plates containing 179 FDA-approved oncology drugs were obtained through the NCI Development Therapeutics Program (DTP). Drugs were provided as 10 mM stock solutions in DMSO. Cells were seeded in 96-well plates (ThermoFisher, Cat. #260887) to reach approximately 30-50% confluency and treated with 1 µM of each drug for 72 hours. Cell viability was assessed using 0.15 mg/mL resazurin (Sigma-Aldrich, Cat. #R7017, prepared as described by Riss et al. [12]) staining and quantified using a Synergy HTX multi- mode reader (Agilent Biotek) with Gen5 software (v5.17). Selected drugs were further evaluated in 24- well plates (Fisher Scientific, Cat. #3422). Cells were seeded to reach approximately 30-50% confluency in 24 hours, then treated with the drugs at varying concentrations. After 72 hours of treatment, the cells were fixed and stained with 0.5% crystal-violet (Gentian, Sigma-Aldrich, Cat. #1004731) in methanol for quantification.

Drug synergy studies were performed by combining compounds at concentrations around their minimum effective doses. Synergy scores were calculated using SynergyFinder Web (version 3.0), utilizing the Zero Interaction Potency (ZIP) Model [13]. A ZIP synergy score <-10 was antagonistic, >10 was synergistic, and between -10 and 10 was additive.

### In vivo drug efficacy studies

To evaluate treatment efficacy for various drugs, mice aged between 6 and 12 weeks were either subcutaneously injected with PDC cells or implanted with PDX tumors and assigned to the following treatment groups. The control group was always categorized as receiving no oncology drug.

For evaluation of romidepsin alone, PDX6 tumors were implanted into the bilateral flanks of mice. No ovariectomy was performed, and the mice were randomized into two groups, control (n=4 mice) and romidepsin (n=3 mice) a few days after implantation. Romidepsin (Bristol Myers Squibb, 0.02 mg/kg in saline, intraperitoneally) was administered three times a week. The mice were sacrificed 21 days after the first treatment.

For evaluation of CUDC-907, 1-2×10^6^ PDC6 or PDC4 cells were injected into the bilateral flanks of mice. No ovariectomy was performed, and for each model system the mice were randomized into two groups: control (n=4 mice for PDX6, n=3 mice for PDX4) and CUDC-907 (n=3 mice for PDX6, n=4 mice for PDX4) two weeks after injection. CUDC-907 (CURIS, 0.02 mg/mouse/100 µL in 30% Captisol, intravenously) was administered three times weekly. The mice were sacrificed 22 days after the first treatment.

For evaluation of romidepsin and azacitidine (5-aza) combination therapy, tumors were implanted into the bilateral flanks of mice from six endometrioid EC xenograft lines. Mice were randomized into two groups for each model: control (n=3 per line) and dual treatment (n=4 for PDX3, PDX50, and PDX57 and n=3 for PDX74, PDX75, and PDX100). Both drugs were administered intraperitoneally at 1 mg/kg; romidepsin weekly and 5-aza three times weekly (Bristol Myers Squibb). Mice were sacrificed between 11 days (PDX3) and 36 days (PDX75) after drug initiation.

Tumor monitoring and euthanasia criteria were assessed as described above. Upon harvest, tumors were imaged, weighed, sectioned for IHC, and snap-frozen for RNA and protein analysis.

### Western blot analysis

Cells were harvested at around 95% confluency, washed with PBS three times, and lysed with lysis buffer (1% NP-40 (RPI, Cat. #1005104), 150mM NaCl, 50mM Tris pH 7.2 (RPI, Cat. #30UC26)) and protease and phosphatase inhibitor cocktail on ice. Lysates were sonicated, centrifuged, and protein concentration was determined using a bicinchoninic acid (BCA) assay. Equal amounts of protein were separated on SDS-PAGE gels, transferred onto nitrocellulose membranes (0.2 μm, Bio-Rad, Cat. #1620112) and blocked with 5% nonfat dry milk in washing buffer (TBST). The membranes were incubated with the primary antibody overnight at 4°C, followed by incubation with appropriate secondary antibodies for 2 hours at room temperature. Antibody details are provided in Supplemental Table 1. Signals were detected with ChemiDoc imaging system (Bio-Rad) and ImageLab, and the membranes were treated using enhanced chemiluminescence (ECL, Clarity Max Western ECL Substrate, Cat. #1705062, Bio-Rad) or SuperSignal West Dura (ThermoFisher, Cat. #34075).

### Statistical Testing and Data Analysis

A Student’s two-sample t-test, assuming unequal variances, was used to test the significance of tumor volume, weight, and patient age in relation to PDX and PDC establishment. The hypothesized mean difference was set to 0, with a significance threshold of α = 0.05. All data analyses were performed in Microsoft Excel, and graphs were generated in both Microsoft Excel and RStudio.

## Results

### Patients’ diagnosis and establishment of corresponding PDX and PDC lines

Between December 2022 and September 2025, we collected 103 matched normal and tumor samples representing all histologic subtypes of EC (Fig. 1A). Endometroid carcinomas comprised the largest subgroup (61.2% of cases), followed by serous carcinomas (15.5% of cases), with additional dedifferentiated, carcinosarcoma, clear cell, squamous, and undifferentiated tumors represented in the cohort (Fig. 1A, D). Using the workflow shown in Fig. 1B, we successfully established 53 PDX models (52.0% engraftment rate) and 15 PDC lines (14.7% of patients; 28.3% of PDXs) (Fig. 1C). PDX establishment rates were similar between low-grade and high-grade tumors overall (51.1% vs 54.0%, p=0.77). However, among endometrioid cases (Grades 1-3, n=63), PDX engraftment success increased with tumor grade, from 42% in grade 1 tumors (n=19) to 57% in grade 2 tumors (n=28) and 63% in grade 3 tumors (n=16) (Fig. 1D). Likewise, recurrent or metastatic high-grade tumors demonstrated higher engraftment rates than their low-grade counterparts (70.0% vs 55.6%), consistent with their greater tumorigenic potential. Among histologic subtypes, dedifferentiated tumors exhibited the highest PDX establishment rate (85.7%), followed by Grade 3 endometrioid carcinomas (62.5%), and carcinosarcomas (60.0%). Recurrent lesions achieved a 100% engraftment rate, whereas metastatic lesions show a 50% engraftment rate. PDC lines were successfully established from all grades of endometrioid, serous, dedifferentiated, and carcinosarcoma subtypes. In addition, successful PDX establishment was associated with older patients (68.6 ± 1.61 n=53 vs. 62.56 ± 1.57 n=50, p=0.004), whereas no association was observed for PDC establishment.

**Fig 1.**
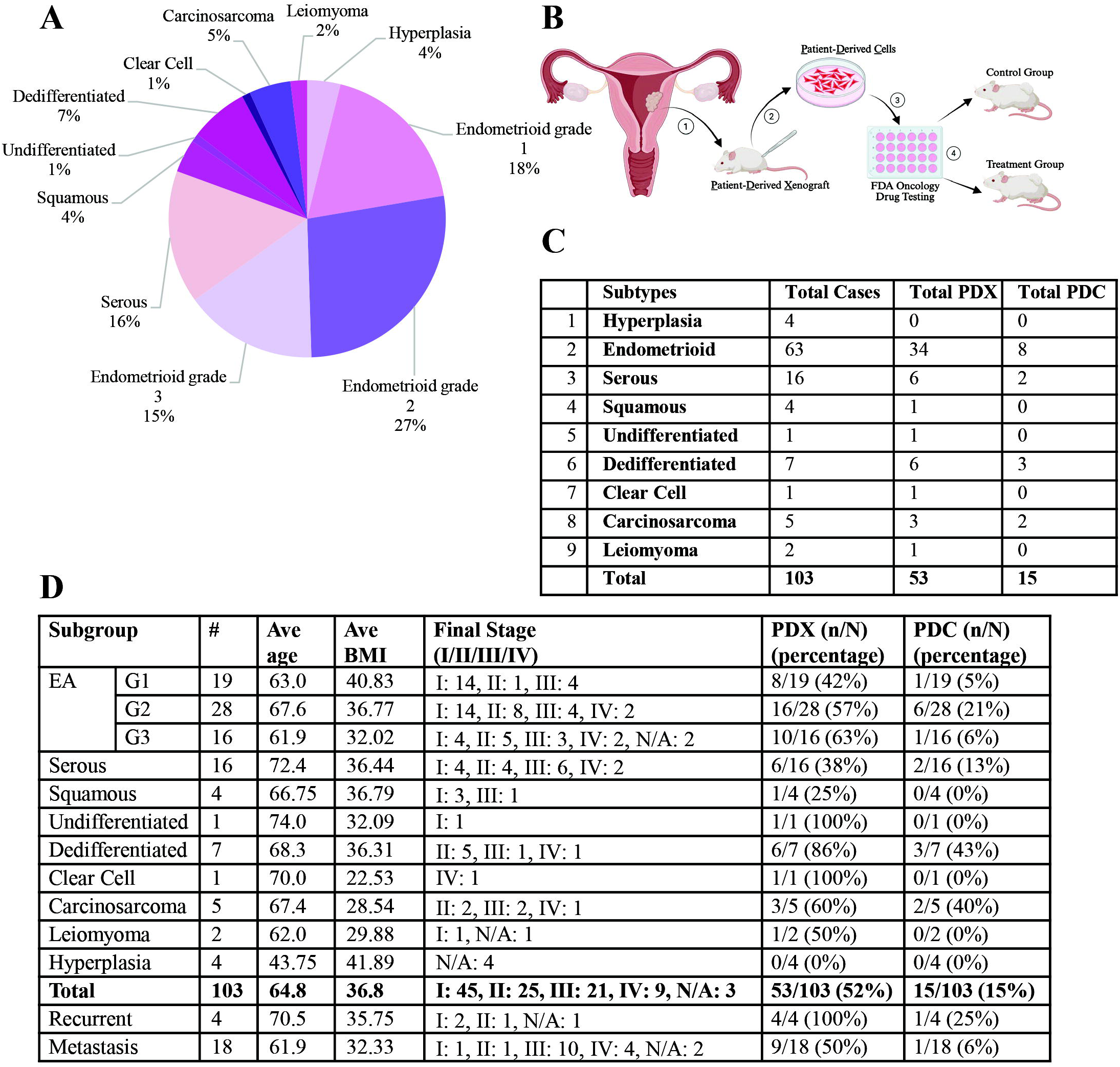
Clinical characteristics and workflow for successful establishment of Patient Derived Xenograft (PDX) and subsequent Patient Derived Primary Cancer Cells (PDC) models. (A) Distribution of endometrial cancer subtypes across patients that we have collected (n=103). Percentages are calculated relative to the total number of cases. (B) Schematic overview of generation of our models and subsequent screening. 1) Establishment of PDX lines: Tumor taken from operating room and implanted in NSG mice. 2) Establishment of PDC lines: The tumor is harvested from the mice and enzymatically digested. 3) Drug screening and dose validation *in vitro*: FDA-approved anticancer drugs are screened on stable cell lines for inhibition of cell proliferation. 4) Validation *in vivo*: The drugs are then tested on NSG mice. (C) Summary of successful establishment of PDX and PDC models from our patients, based off diagnosis. (D) Clinicopathologic characteristics of our patients, including average age, body mass index (BMI), tumor stage, and success rate of PDX and PDC establishment.

### Tumor Gene Mutation Analysis

Mutation analysis was performed on tumor samples from selected patients (Fig. 2). The mutational landscape revealed substantial intertumoral heterogeneity. Recurrent alterations were identified in multiple pathways commonly implicated in EC, including chromatin remodeling/epigenetic regulation, PI3K-AKT-mTOR signaling, RTK-MAPK signaling, DNA damage response/genome integrity, and TP53/cell-cycle control pathways (Fig. 2A). The most frequently altered genes included ARID1A (52%), PTEN (42%), TP53 (39%), PIK3CA (29%), KMT2D (23%), and KRAS (16%) (Fig. 2A).

**Fig 2.**
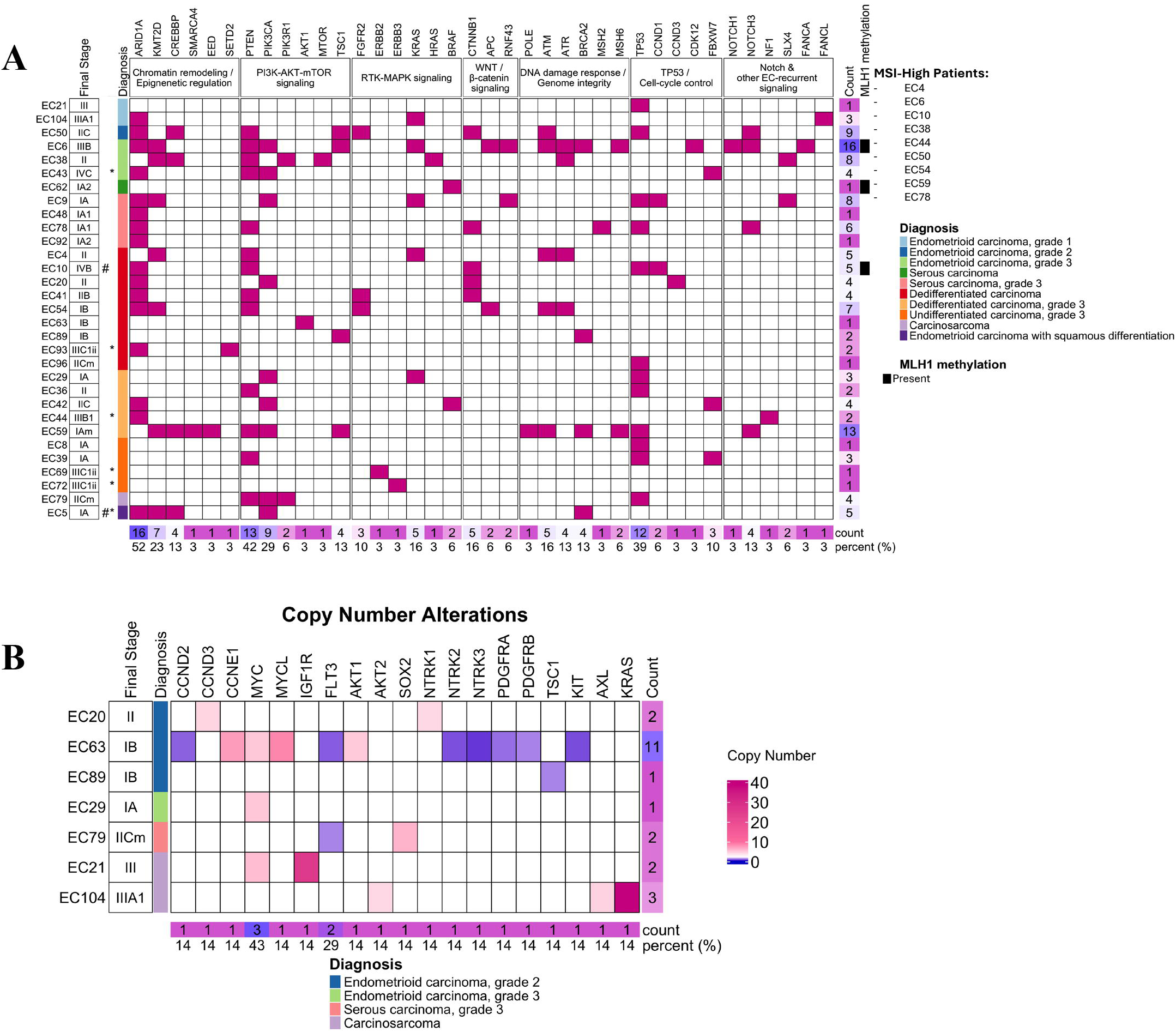
Targeted sequencing on EC-PDX tumors characterizes recurrent somatic mutations in key endometrial cancer–associated genes, showing intertumoral heterogeneity. (A) Depiction of the presence and type of mutations across individual PDX models. * = recurrent, # = metastasis. (B) Copy number alterations, seen in a seven-patient subset. A copy number of two is considered normal. Frequently altered copy numbers include *MYC*, which showed copy number increases, and *IGF1R*, which showed copy number decreases. This genomic diversity provides a molecular rationale for differential drug sensitivity observed across PDX and derived PDC models.

Overall, MLH1 methylation was detected in 18 cases (3 concurrent with mutation data), suggesting potential mismatch repair deficiency (MMR-d). Mutation burden varied substantially between tumors, ranging from 1 detectable mutation in EC8 and EC13 to 16 mutations in EC6 (Fig. 2A). Copy number analysis identified localized amplification in selected tumors, including IGF1R (25 copies) and MYC (5.4 copies) in EC21(Fig. 2B). Several cases (EC44, EC48, EC50, and EC54) did not yield reportable mutation data.

### Establishment and characterization of PDX models

Successful engraftment of human tumors in NSG mice was confirmed by GFP imaging and microscopic analysis (Supple. Fig. 1A). Histologic evaluation demonstrated that PDX tumors largely preserved the morphologic features of their corresponding patient tumors across serial passages (Fig. 3A). H&E staining of matched normal tissue, primary tumors, and PDX tumors (F0–F1) revealed strong histologic concordance, supporting the fidelity of the PDX models.

**Fig 3.**
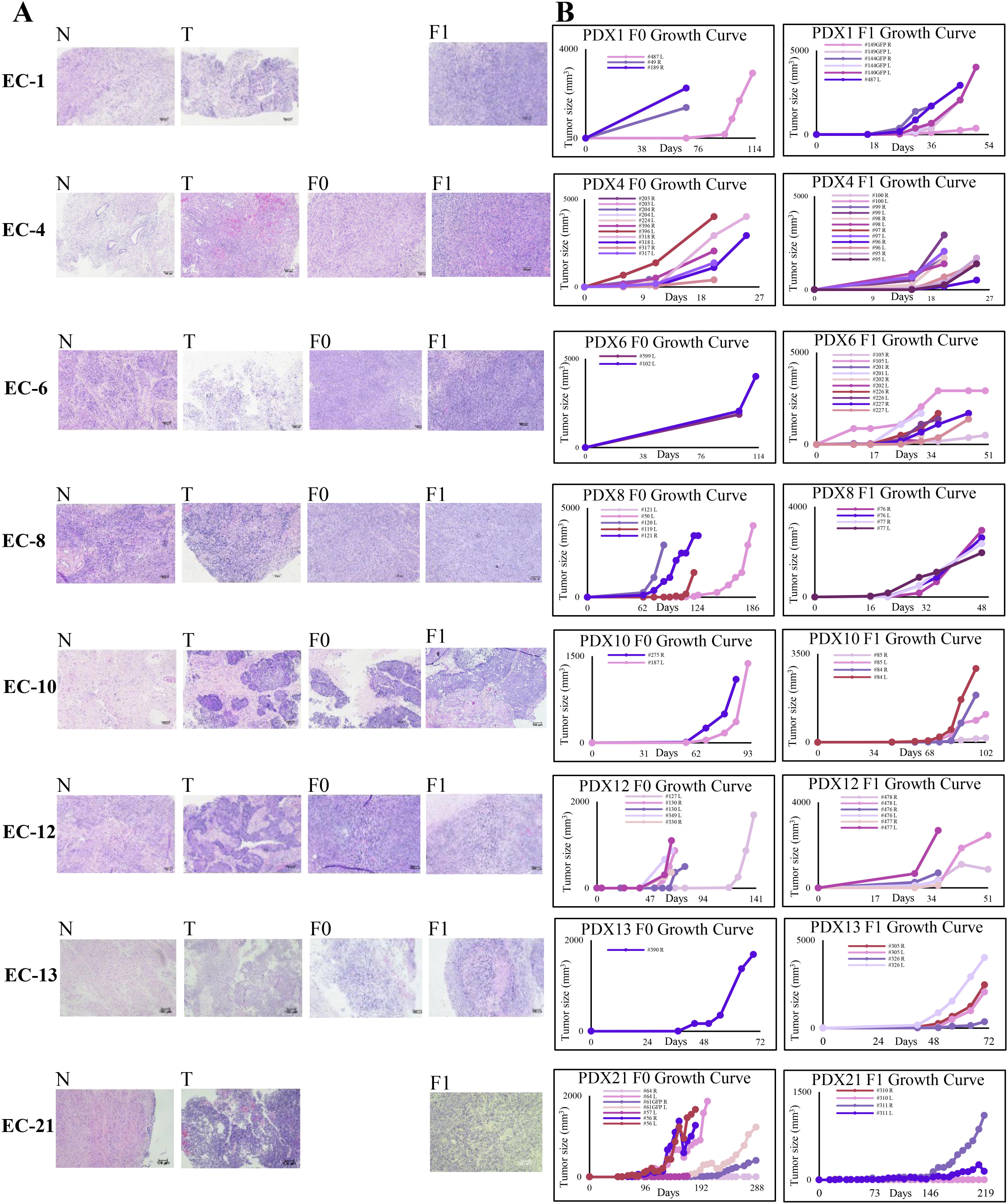
PDX models match patient tumor histology while displaying heterogenous growth dynamics between patients. (A) Hematoxylin and eosin (H&E) stains for each human tumor of our stable cell lines. N is normal human tissue, T is the human tumor, F0 is the initial generation in mice, and F1 is the second generation in mice. Histopathologic features are maintained between patients and mice. (B) Tumor volume charts for each PDX. The # represents the mouse ID number, “R” indicates right side tumor, “L” indicates left side tumor.

Tumor growth kinetics varied substantially across PDX models (Fig. 3B). Tumor initiation occurred within 1–8 weeks (median, 3 weeks), whereas progression to 20 mm ranged from 8–20 weeks (median, 12 weeks). Variations in tumor growth rates are observed with tumors derived from the same patient but implanted on different flanks, suggesting that tumor microenvironmental factors or individual differences may influence tumor progression (Fig. 3B).

### Establishment and characterization of PDC models

Fifteen PDC lines were successfully established from endometrioid, serous, dedifferentiated, and carcinosarcoma tumors (Fig. 4A). Representative brightfield images demonstrated substantial heterogeneity in cellular morphology, growth pattern, and proliferative capacity (Fig. 4B). Some PDCs displayed predominantly adherent epithelial-like growth (PDC1 and PDC10), whereas others formed floating aggregates or spheroid-like clusters (PDC6, PDC12, PDC54 and PDC96). Doubling times ranged from 21 hours (PDC4) to 120 hours (PDC96). Most PDCs display an average doubling time of 36 hours, indicating that they are stable and viable for *in vitro* expansion (Fig. 4B).

**Fig 4.**
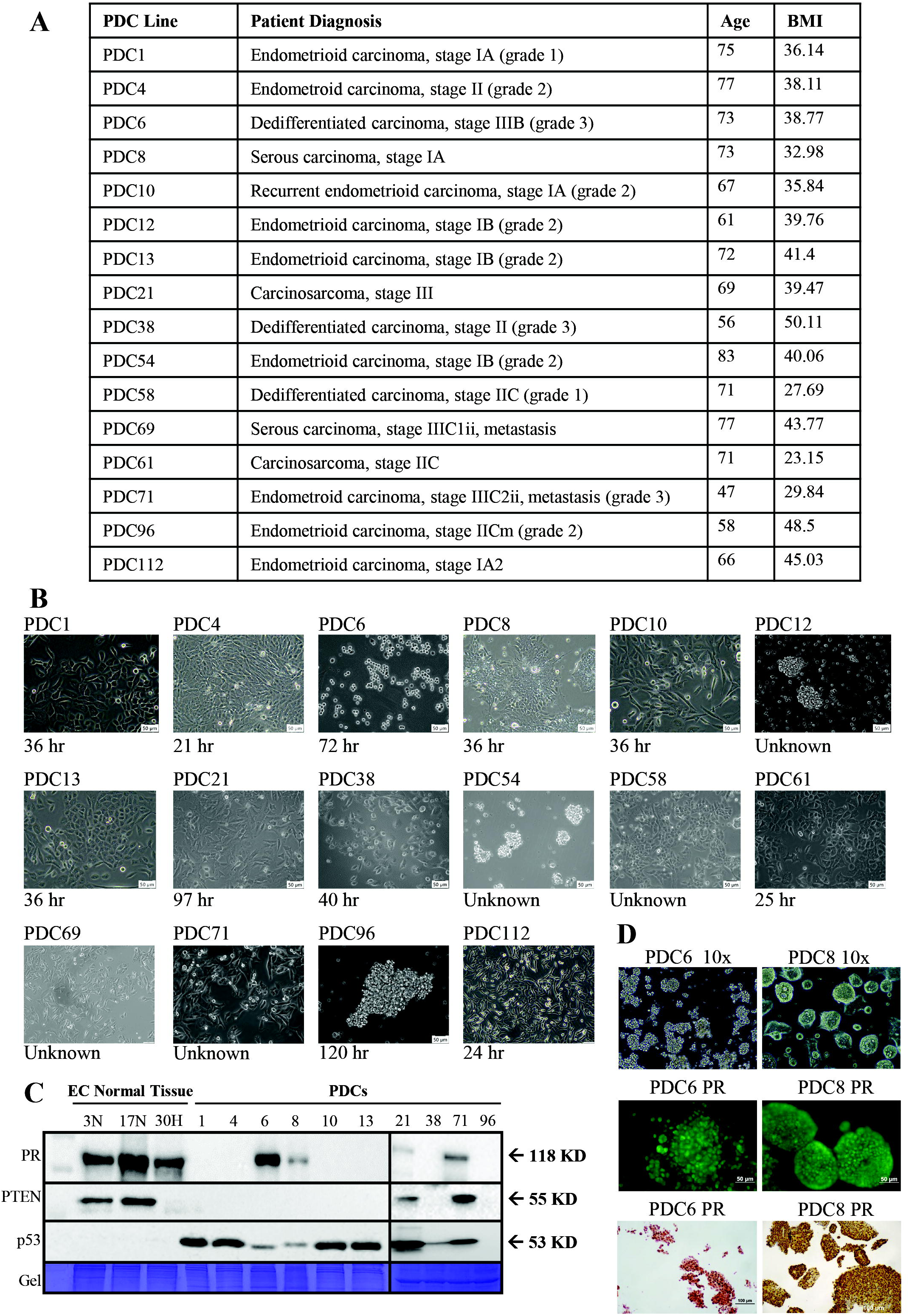
Establishment and phenotypic characterization of novel endometrial cancer PDC models. (A) Clinical characteristics for the patients for each cell line created. (B) Representative brightfield PDC images. The doubling time of each PDC line is indicated below the images, demonstrating variability in proliferative capacity across patient-derived samples. Scale bars = [50 μm]. (C) Western blotting for stable PDC lines, compared to normal tissue (N) and hyperplasia (H), showing upregulated p53 which is characteristic of oncogenic tissue, along with occasional expression of PR which is important for therapeutic response. The expression patterns vary across models, continuing to reflect tumor heterogeneity. (D) Spheroid formation, immunofluorescence (IF) analysis, and IHC for PR expression of PDC6 and PDC8, showing that both cell lines are PR positive.

Western blot analysis further demonstrated molecular heterogeneity among PDC models (Fig. 4C). Compared with normal endometrial tissue and hyperplasia controls, all evaluated PDC lines exhibited elevated p53 expression, which is consistent with oncogenic tissue. In contrast, PR and PTEN expression varied considerably among models, reflecting the biological diversity of the original patient tumors.

PDC6 and PDC8 cells successfully formed spheroids in 3D culture (Fig. 4D). Under low-magnification microscopy, PDC6 and PDC8 spheroids appear compact with some irregular morphology. Immunofluorescence labeling reveals progesterone receptor (PR) expression within spheroids, with a higher signal intensity in PDC8.

### High-Throughput Drug Screening on PDCs

High-throughput drug screening was performed using 13 PDC models with a library of 179 FDA-approved oncology drugs to assess growth-inhibitory effects of various targeted and chemotherapeutic agents (Fig. 5). Screening with NCCN-recommended agents revealed limited efficacy across many models, highlighting substantial variability in therapeutic response (Fig. 5A, 5B, and Supple. Fig 2).

**Fig 5.**
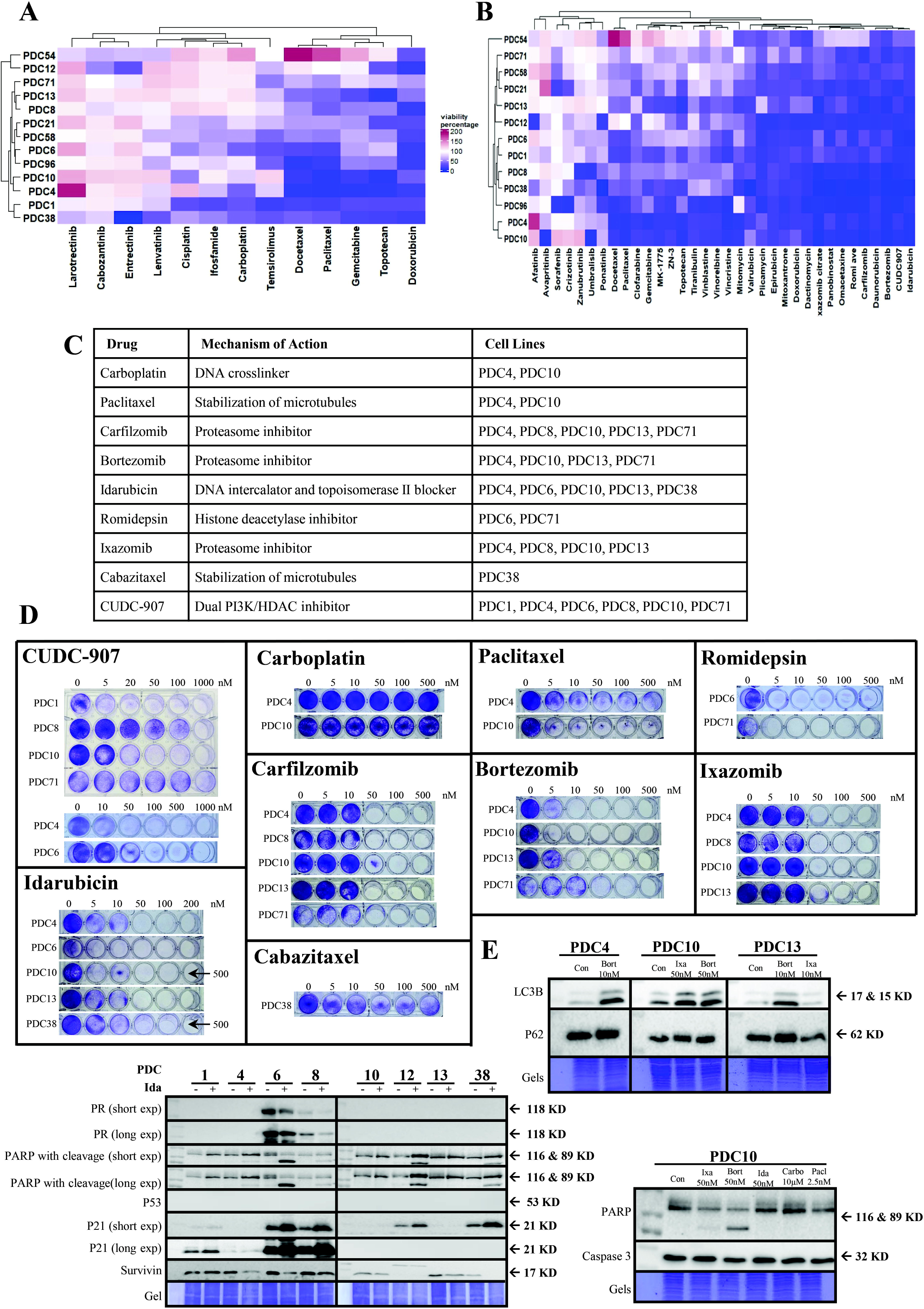
Screening of FDA approved oncology drugs identifies therapeutic variabilities in PDC models. (A-B) IC90 analysis of FDA-approved and selected drugs on PDCs. 100% viability percentage is shown with white and indicates no change compared with control. (A) IC90 analysis of NCCN guideline drugs for EC treatment on stable PDC lines. (B) IC90 drug sensitivity heatmap of treatment of stable PDCs with 179 FDA-approved and other selected drugs. The most potent inhibitors (>90% growth suppression) include idarubicin, the dual PI3K/Akt inhibitor CUDC-907, and two proteasome inhibitors, bortezomib and carfilzomib, with variable responses across PDC lines. (C) Table of most effective drugs identified in IC90 screening, including carboplatin and paclitaxel which are considered the standard-of-care drugs for EC. Their mechanisms of action and the cell lines that they were tested on are also shown. (D) Dose response validation of selected drugs in stable cell lines. (E) Western blot analysis of drug treatment on different PDC lines. PDC4, PDC10, and PDC13 were treated with various FDA oncology drugs, and eight PDCs were treated with idarubicin (Ida). Key autophagy indicators like LC3B cleavage were seen throughout, but PARP cleavage only occurred in a few PDC lines. Doses were determined based on dose-response experiments in (D).

Consistent with the heterogeneous nature of EC, the broader FDA-approved drug library identified marked differences in drug sensitivity among PDCs.

Several compounds demonstrated potent growth-inhibitory activity across multiple models, including HDAC inhibitor romidepsin, dual PI3K/HDAC inhibitor CUDC-907, anthracycline idarubicin, and proteasome inhibitors bortezomib, carfilzomib, and ixazomib (Fig. 5B, C).

Well plate assays after drug treatment further validate these findings (Fig. 5D). Trends are consistent with the inhibition rates shown in the heatmaps, further confirming the reliability of the high-throughput screening results. Western blot analyses revealed drug specific molecular effects across PDC lines. Idarubicin induced PR expression in PDC6 and PDC8, while proteasome inhibitors bortezomib and ixazomib induced PARP1 cleavage in some cell lines, and LC3B cleavage in all.

Idarubicin also increased p21 expression in multiple PDCs. (Fig. 5E).

To determine whether drug responses observed in 2D cultures were preserved in a three-dimensional setting, PDC54 spheroids were treated with selected candidate drugs. Romidepsin, CUDC-907, mitoxantrone, and idarubicin each reduced spheroid growth (Supple. Fig. 1B), consistent with their inhibitory effects in matched 2D cultures. These findings suggest that 2D PDC models provide a reliable platform for high-throughput drug screening.

### Synergistic Effects on our PDC Models

The antiproliferative activity of romidepsin with mitoxantrone or idarubicin was assessed across different concentration combinations, and synergy heatmaps and 3D surface plots were generated to visualize interaction patterns (Supple. Fig. 3). The results showed additive synergistic interactions between romidepsin and idarubicin within specific concentration ranges, with ZIP synergy scores above 0 for all PDCs, and above 10 in PDC1 and PDC6. Similar effects were observed between romidepsin and mitoxantrone, where ZIP synergy scores are above 0 for all PDCs and PDC1, PDC6, and PDC8 exceed 10. The strongest effects were observed in the 3D surface plots, where peak synergy regions vary based on drug dose and cell line. Together, these findings suggest that romidepsin can enhance the antitumor activity of both mitoxantrone and idarubicin in select PDC models.

### Drug efficacy in NSG mice

Five individual drugs and two combination therapies were evaluated *in vivo* after showing significant success *in vitro*. All drugs tested reduced tumor growth to varying degrees.

Romidepsin and 5-aza combination therapy significantly reduced tumor volume in all PDX lines, and terminal tumor weight in 5 of 6 PDX lines (Fig. 6A-E). One PDX75 mouse was found dead 11 days after drug treatment despite showing healthy weight and no adverse effects to the drugs. Further IHC analysis of tumors after extraction showed elevated PR expression in tumors treated with the dual therapy compared with the control (Fig. 6D).

**Fig 6.**
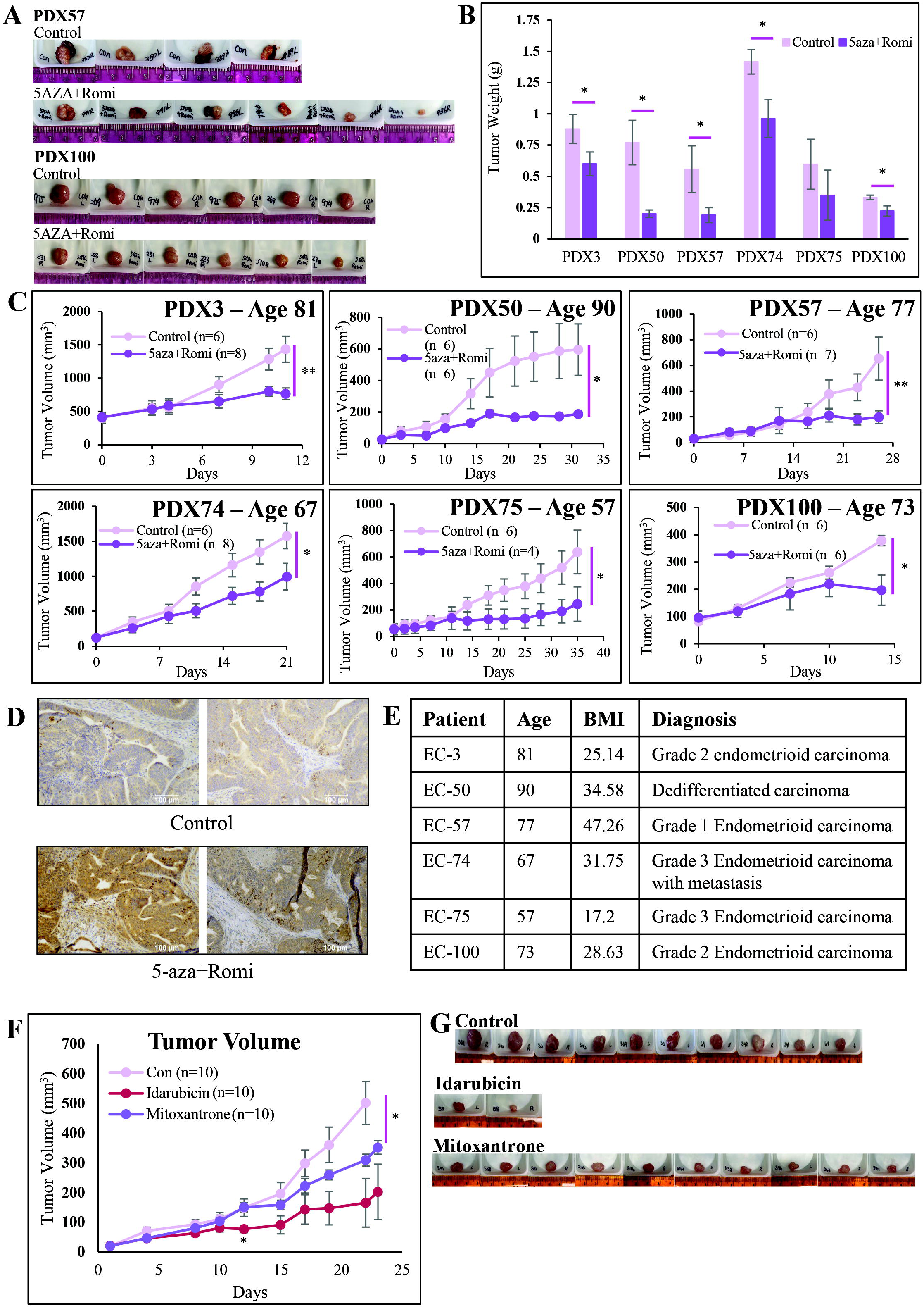
*In vivo* validation of candidate therapies demonstrates efficacy in representative PDX models. (A-E) Evaluation of dual epigenetic therapy with 5-azacitidine (5-aza) and romidepsin (Romi) across 6 PDX models. These patients are on the higher age range for endometrial cancer. (A) Images of representative tumors from PDX57 and PDX100. (B) Tumor weight charts for these PDX lines, comparing control with 5-aza and Romidepsin dual-therapy. Significance is as follows: PDX3 p=0.044, PDX50 p=0.013, PDX57 p=0.045, PDX74 p=0.014, PDX75 p=0.19, and PDX100 p=0.04. (C) Tumor volume charts for each PDX line tested. Significance is as follows: PDX3 p=0.008, PDX50 p=0.027, PDX57 p=0.006, PDX74 p=0.021, PDX75 p=0.049, PDX100 p=0.019. (D) IHC data for PDX100, stained for PR. (E) Table summarizing patients used for this evaluation. (F-G) Evaluation of chemotherapeutic devices idarubicin and mitoxantrone. (F) Tumor volume graph for idarubicin and mitoxantrone. Mitoxantrone significantly reduced tumor volume compared to the control through the experiment (p=0.037). Idarubicin had significantly lower tumor volume 12 days after initial drug treatment than the control (p=0.021). (G) Images of representative tumors from the mice following harvest. Data is presented as mean ± SEM. Statistical significance is indicated as *p < 0.05, **p < 0.01.

Mitoxantrone also significantly reduced tumor volume in mice compared with control. Idarubicin demonstrated significant antitumor activity but was associated with substantial toxicity in mice (Fig. 6F, G). A significant reduction in tumor volume was observed 12 days after treatment initiation, when all mice remained alive. Since both idarubicin and mitoxantrone tumors were extracted prior to the control, terminal tumor weights were not available for direct comparison.

Romidepsin and CUDC-907 both reduced tumor weight and volume on average, but these effects were not statistically significant (Supple. Fig. 4). Western blot analysis reveals decreased SETDB1 and p- Akt and increased H3Ace when treated with CUDC-907 (Supple. Fig. 4F, I).

## Discussion

Endometrial cancer exhibits substantial histologic, molecular, and therapeutic heterogeneity, creating a major challenge for the development and evaluation of effective therapies [14]. In this study, we established an integrated PDX and PDC platform representing all histologic subtypes of EC and demonstrated that these models faithfully preserve patient tumor characteristics while capturing diverse treatment responses [7, 8, 15]. Functional screening of 179 FDA-approved oncology drugs identified considerable variability in drug sensitivity and revealed several unexpected therapeutic vulnerabilities, including proteasome inhibition, dual PI3K/HDAC inhibition, and epigenetic combination therapy. To our knowledge, this represents one of the largest integrated EC PDX and PDC platforms reported to date. Together, these findings establish a valuable preclinical resource for therapeutic discovery, biological investigation, and translational studies in EC. Unlike previous EC PDX studies that primarily focused on model establishment or molecular characterization, our study integrates patient-derived xenografts with matched primary cancer cell models, large-scale functional drug screening, and *in vivo* validation, providing a comprehensive platform for therapeutic discovery. Our platform achieved a PDX establishment rate comparable to or slightly higher than those reported previously (Fig. 1C) [10]. Historically, limitations with PDX models include prolonged establishment time and low primary engraftment success rate, especially in early-stage cancers [16, 17]. Our PDX platform helps address these challenges by both reducing time for establishment and increasing the success rates across multiple stages and histologic subtypes of EC. In addition, we successfully established PDX lines from all EC subtypes. Several technical factors have likely contributed to the successful establishment of our PDX platform. Unlike conventional approaches, freshly resected patient tumors were sectioned into thin fragments prior to implantation, thereby improving nutrient diffusion and reducing ischemic injury. The site of tumor implantation is especially important in the engraftment rate [18]. We chose to implant tumors in highly vascularized flank regions of NSG mice, which may facilitate early tumor engraftment and growth.

Patient-derived cell lines (PDCs) are an extension of PDX technology, offering significant advantages in speed, scalability, and cost efficiency. In successful cases, PDCs can be established and ready for experimental use within just 2 to 7 weeks (Fig. 4B). PDCs are particularly well suited for large scale, high-throughput drug screening (Fig. 5). Compared to 3D organoid models, 2D PDC lines are easier to culture and standardize [16]. In our study, both 2D and 3D culture systems demonstrated similar responses to drug treatment, validating the use of 2D cell lines and supporting the use of conventional 2D PDC cultures for efficient functional drug screening. The genomic profiles of our models closely mirrored the known molecular landscape of EC, with recurrent alterations in ARID1A, PTEN, TP53, PIK3CA, KRAS, and KMT2D [19]. These findings support the biological relevance of the platform and provide opportunities for future genotype-directed therapeutic studies (Fig. 2A) [20, 21].

Establishment of PR-positive models has been challenging, with relatively few reported in the literature. In 2022, Rush et al. were able to establish cell line HCI-EC-23 as an estrogen and progesterone responsive cancer cell line for EC [22]. We successfully established two PR-positive cell lines, PDC6 and PDC8. Imaging PDC lines revealed variability in growth patterns and cellular morphology, corresponding to the diagnoses of patients (Fig. 4B). Western blot analysis further confirmed relevant oncogenic signaling, including consistent p53 and variable PR and PTEN expression (Fig. 4C) [23].

When tested with chemotherapeutic agents, our stabilized cell lines showed minimal responsiveness to most of the NCCN Guideline drugs for EC, where all drugs besides doxorubicin increased the viability of cells in at least one of our PDCs (Fig. 5A) [24, 25]. The standard-of-care chemotherapy carboplatin did not decrease cell proliferation at high doses, and paclitaxel exhibited a cytotoxic plateau [26].

Screening with FDA-approved and selected drugs demonstrated superior activity with idarubicin (topoisomerase inhibitor), CUDC-907 (dual PI3K/HDACi), and bortezomib (proteasome inhibitor) showing consistent reduction of cell proliferation across all PDCs tested (Fig. 5B). Romidepsin (HDACi), mitoxantrone (topoisomerase inhibitor), ixazomib (proteasome inhibitor), and carfilzomib (proteasome inhibitor) were also effective in large-scale screening. Subsequent dose response demonstrated potent activity at low nanomolar concentrations, with idarubicin and bortezomib reducing cell proliferation at concentrations of 10 nM and below (Fig. 5D). Western blot mechanistic analysis revealed drug-specific apoptotic and autophagic molecular effects (Fig. 5E). Bortezomib and ixazomib resulted in LC3B cleavage into LC3-I and LC3-II, which is a key indication of autophagy [27, 28]. These drugs are FDA-approved for multiple myeloma and mantle cell lymphoma and are actively used in over 100 clinical trials according to clinicaltrials.gov [29–31]. Idarubicin led to PARP1 cleavage in 5 of 8 PDC models tested, suggesting apoptosis induction [32, 33].

Chemotherapeutic agents were tested on the *in vivo* level, with either the implantation or subcutaneous injection of tumors into NSG mice. Mitoxantrone, idarubicin, CUDC-907, romidepsin, and romidepsin with 5-aza all reduced tumor size *in vivo* (Fig. 6 and Supple. Fig. 4). However, idarubicin exhibited significant systemic toxicity, leading to early mortality in most treated mice. The same effects were observed in healthy male mice. The drug is actively tested in clinical trials but is administered in a liposomal form. Using this nano-formulation reduces the toxicity of anthracycline antibiotics like idarubicin [34]. Idarubicin hydrochloride is actively being investigated in over 60 clinical trials for effectiveness in a variety of cancer types, and is FDA approved for acute myeloma leukemia [31, 35]. Romidepsin and 5-aza proved effective in all PDX models tested. Age-associated DNA methylation changes have been implicated in endometrial cancer progression and may contribute to loss of hormone responsiveness in postmenopausal patients [36]. As a DNA methyltransferase inhibitor, 5-aza reverses aberrant DNA methylation. Its deoxy analog, 5-aza-2’ deoxycytidine has previously demonstrated activity in EC [37]. Our patients used for this analysis were 74 years old on average, and we even found an increase in PR between control and treatment groups in PDX100 (Fig. 6C, D).

Romidepsin and 5-aza have been used in 5 clinical trials, 2 of which are still active, for cancers like lymphoma, especially peripheral T-cell lymphoma (PTCL) [38, 39]. PTCL and EC have similar characteristics in terms of reliance on genetic and hormonal characteristics and share responsiveness to drugs like doxorubicin in combination with other therapies [40, 41]. In our study, combined inhibition of DNA methylation and histone deacetylation restored PR expression and simultaneously suppressed tumor growth.

There are several limitations to our studies. Our work is limited to the small number of established PDC lines. Though we were able to establish 15 cell lines for drug screening, we are lacking lines representing major subtypes including squamous and undifferentiated. Our PDC library would be strengthened with the inclusion of all subtypes for EC. Future studies will incorporate hormonal and immunotherapies into our testing, to continue validating our platform for use against EC. Our *in vivo* studies rely on NSG mice, which are immunodeficient to keep them from rejecting the implanted human tumors. Comprehensive molecular profiling was not available for all patient tumors. Therefore, correlations between specific genomic alterations and therapeutic responses could not be systematically evaluated. Drug selection was based primarily on functional screening rather than mutation-guided therapeutic matching. Future studies leveraging the existing PDX/PDC biobank may allow systematic evaluation of genotype-drug response relationships, including tumors harboring PIK3CA mutations, TP53 alterations, ARID1A loss, or oncogene amplifications.

Consistent with previous reports, therapeutic responses and growth characteristics varied substantially among EC subtypes [42] These observations further emphasize the need for diverse, well-characterized patient-derived model systems capable of capturing this biological heterogeneity. As our PDX/PDC biobank continues to expand, it will provide an increasingly valuable resource for mechanistic studies, therapeutic evaluation, and clinical translation approaches in endometrial cancer.

## Conclusions

In conclusion, we established a comprehensive endometrial cancer PDX and PDC platform spanning multiple histologic subtypes and molecular backgrounds. This resource faithfully recapitulates patient tumor biology, supports large-scale functional drug screening, and enables *in vivo* therapeutic validation. Application of this platform identified several promising therapeutic candidates and highlighted combined romidepsin and 5-azacytidine treatment as a potentially effective strategy for endometrial cancer. Collectively, this integrated PDX/PDC platform provides a valuable translational resource for precision oncology and future therapeutic development in EC.

## Abbreviations

EC: Endometrial cancer
PDX: Patient-derived xenograft
PDC: Patient-derived primary cancer cell line
PI3K: phosphoinositide 3-kinase
HDACi: Histone Deacetylase inhibitor
5aza: 5-azacytidine
Romi: romidepsin
Ida: idarubicin hydrochloride
Mito: mitoxantrone
Ixa: ixazomib citrate
Bort: bortezomib
Carf.: and carfilzomib

## Data Availability

This data is available upon request.

## CRediT Authorship Contribution Statement

**Tianyue Li**: Data acquisition, Formal analysis, Investigation, **Fang Huang**: Data acquisition, Formal analysis, Investigation, Writing – original draft, **Xiaohao Huang**: Data acquisition, Formal analysis, Investigation, **Ella Pate**: Data Acquisition, Formal Analysis, Investigation, Writing – Secondary draft including full discussion and large alterations to methods and results, **Riley Rosenmeyer**: Data Acquisition, Investigation, **Mya Messenger**: Data Acquisition, Investigation, **Kaylee McSweeney**: Writing – secondary draft, **Samuel Robinson**: Data acquisition, **Adam Deters**: Data acquisition, **Lauren Buchanan**: Data acquisition, **Maggie Meehan**: Data acquisition, Investigation, **Nandini Patel**: Data acquisition, **Allison Diekema**: Data acquisition, **Yiqin Xiong**: Data acquisition, **Xiaofang Zhang**: Data acquisition, **Xiangbing Meng**: Data acquisition, **Shujie Yang**: Conceptualization, Funding acquisition, Investigation, Methodology, Project administration, Supervision, Writing – original draft, Writing – review & editing.

## Acknowledgements

This project was funded through NIH R37CA238274 (SY), U01CA272424 (BL, SY) Administrative Supplements Award, the Department of Pathology Start-Up Fund (SY), the Cancer Research Opportunities at Iowa program (NCI R25 YES Research Education Program 1R25CA273964), the American Cancer Society DICR INTR-24-1297273-01-DICR INTR Internship award, and the Holden Comprehensive Cancer Center at The University of Iowa and its National Cancer Institute Award P30CA086862. We would also like to thank the Comparative Pathology Laboratory (CPL) lab in the Department of Pathology at the University of Iowa for all embedding, IHC, and H&E staining.

## Declaration of generative AI and AI-assisted technologies in the manuscript preparation process

During the preparation of this work, the authors used Microsoft Copilot for checking spelling and grammar. The authors reviewed and edited the output as needed and take full responsibility for the content of the published article.

