## Supplemental Material for "Integrated Patient-Derived Xenograft and Patient-Derived Cell Models Reveal Therapeutic Vulnerabilities Beyond Standard-of-Care Therapy in Endometrial Cancer"

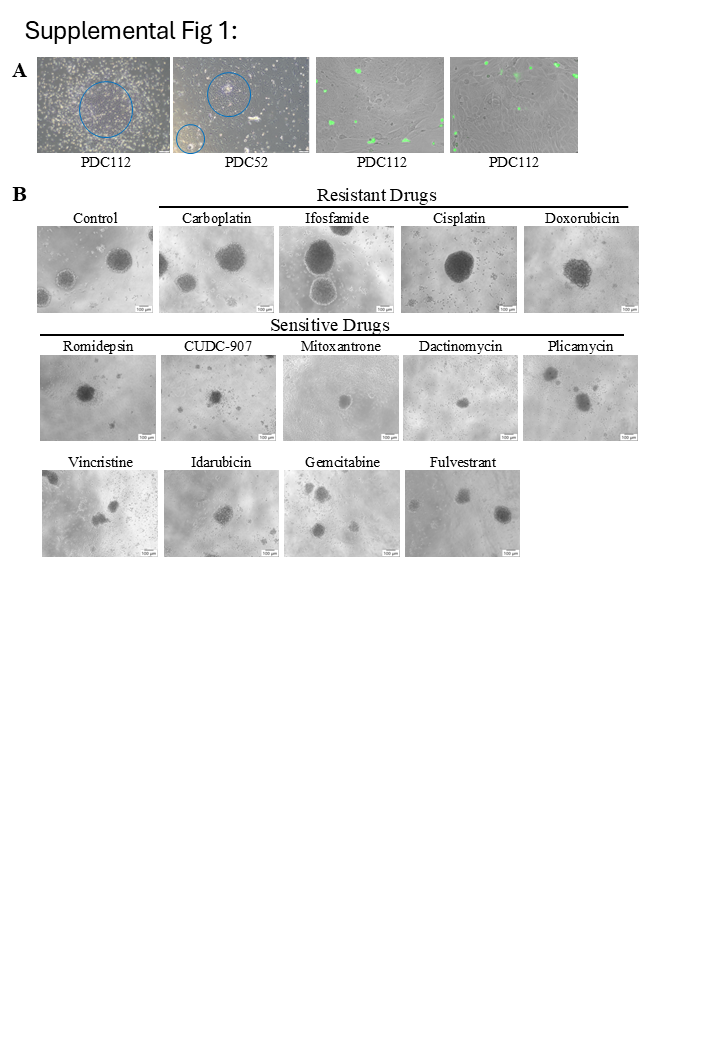

**Supplemental Fig 1.** Establishment and characterization of PDC models.

(A) Microscopic visualization of tumor-derived cells following harvest and digestion of human tumors extracted from mice. In the first two images, human tumor cells are indicated by blue circles, surrounded by normal mouse tissue. In the second two images, GFP labeling identifies mouse cells, while the surrounding tissue that lacks the GFP signal are human tumor cells. (B) Drug response of PDC54 spheroids following treatment with selected FDA drugs. These images demonstrate the effects of drugs on 3D spheroid cultures.

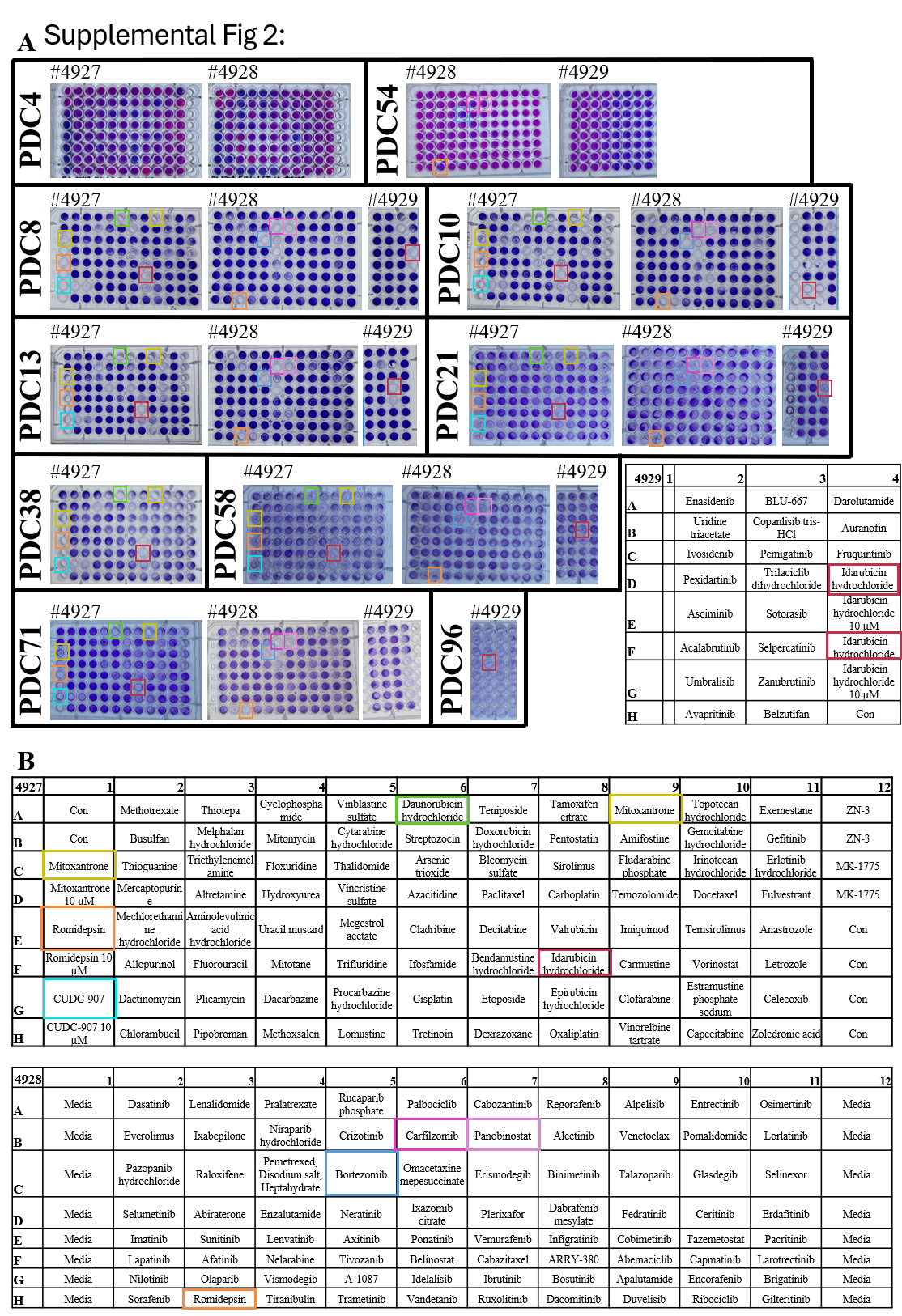

**Supplemental Fig 2.** High-throughput screening of 179 FDA-approved and selected oncology drugs.

(A) Representative 96-well plate assays across several PDC lines. PDC4 and PDC54 were stained with resazurin, where darker blue indicates higher drug efficacy. All other plates were stained with crystal violet, where less purple indicates higher drug efficacy. (B) Plate maps showing the location of each drug. Drugs were diluted to 1 µM unless otherwise noted. Plates 4927 and 4928 were always seeded the same, but plate 4929 showed occasional variation. Boxing for idarubicin hydrochloride is correct based on the map for that plate. Drugs that reduced average cell confluency to less than 15% on average are boxed.

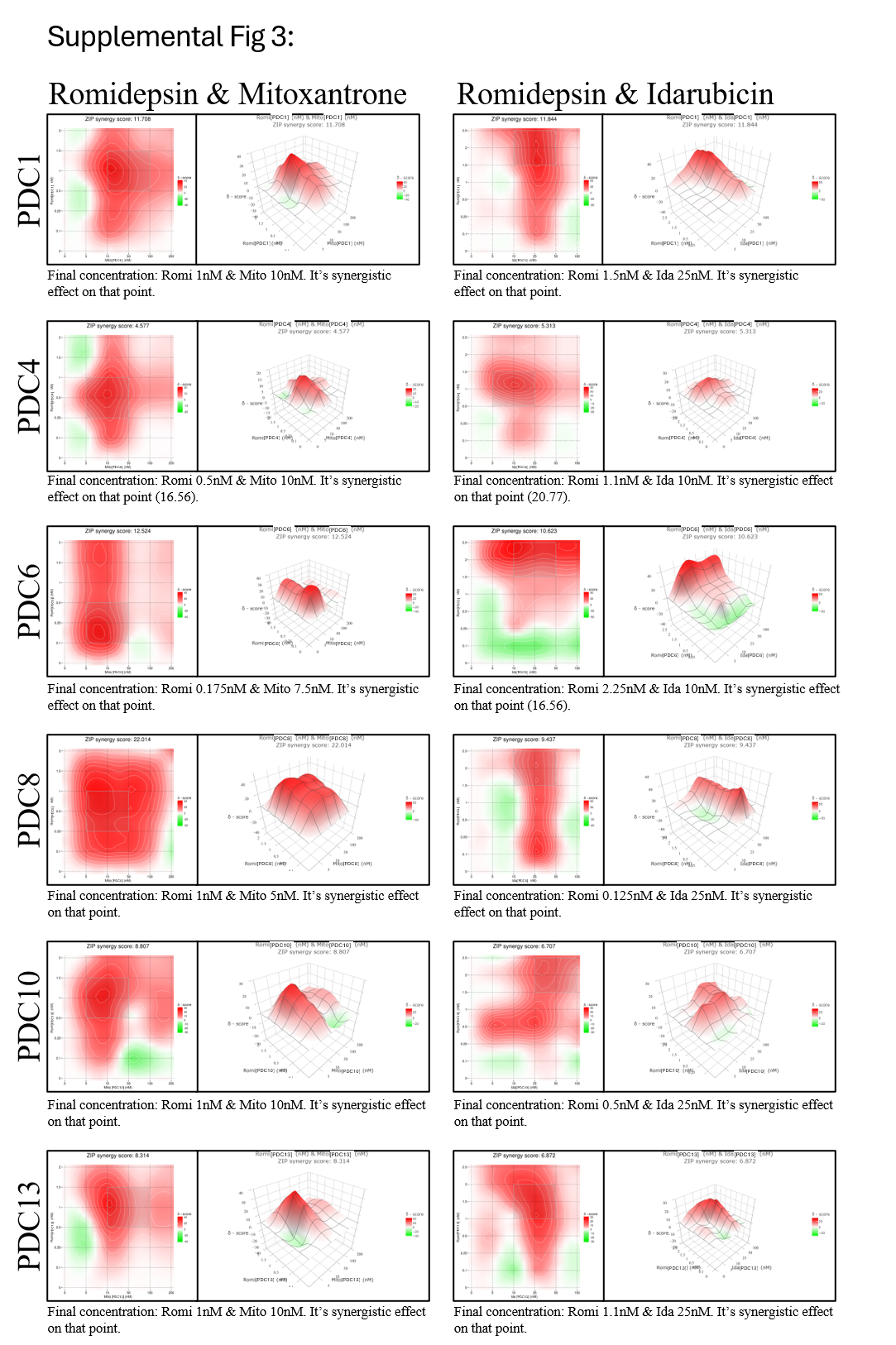

**Supplemental Fig 3.** Synergistic analysis comparing mitoxantrone (Mito) and idarubicin (Ida) to romidepsin (Romi) in PDC models.

This relies on the Zero Interaction Potency (ZIP) Model. Synergy scores can be interpreted as follows: scores < -10 indicate antagonist interactions, scores between -10 and 10 indicate additive effects, and scores > 10 indicate synergistic interactions. Analyses were run using SynergyFinder (web version 3), and the curves were fitted using a four-parameter logistic (LL4) model. The left 2D contour plot and the right 3D surface plot represent the ZIP synergy scores for different drug concentration combinations, with color intensity indicating the strength of the synergy effect. Zoom in for better visualization of the labels.

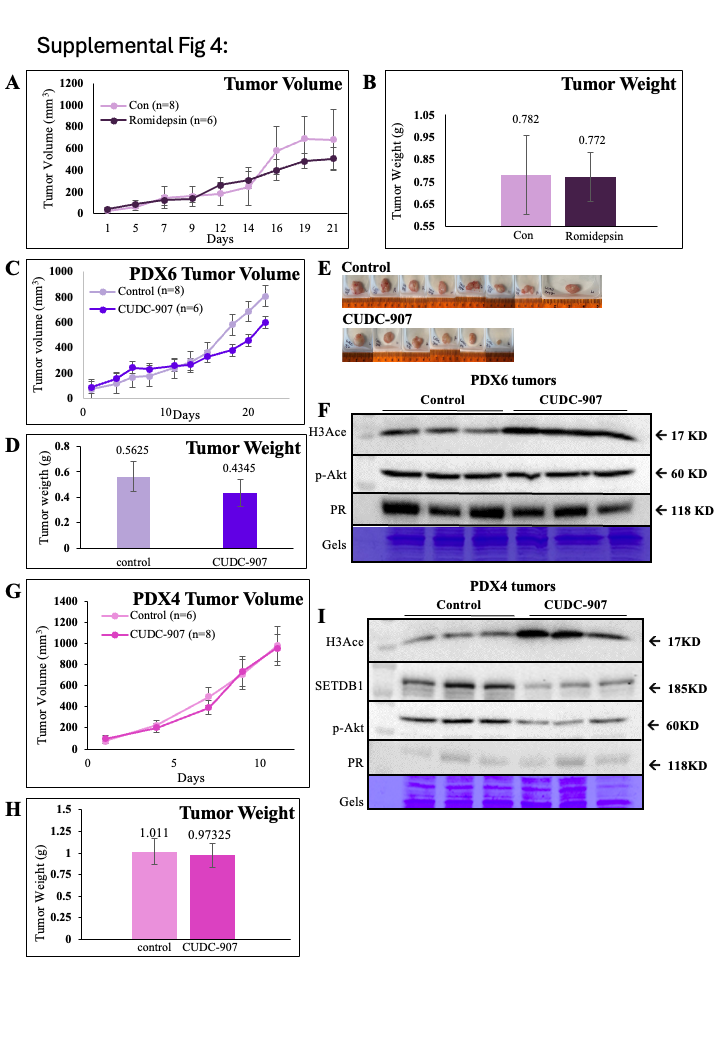

**Supplemental Fig 4.** Additional *in vivo* validation.

(A-B) Validation of romidepsin in PDX6. (A) Tumor volume graph. (B) Tumor weight chart. (C-F) Validation of CUDC-907 in PDX6. (C) Tumor volume graph. (D) Tumor weight graph. (E) Images of representative tumors from the mice following harvest. (F) Western blot comparing the tumors in the control mice to the treatment group. Increased H3Ace was consistent throughout. (G-I) CUDC-907 validation in PDX5. (G) Tumor volume graph. (H) Tumor weight chart. (I) Western blot comparing the tumors of mice in the control and treatment groups. H3Ace is upregulated throughout, while SETDB1 and p-Akt are downregulated. Data is presented as mean ± SEM.

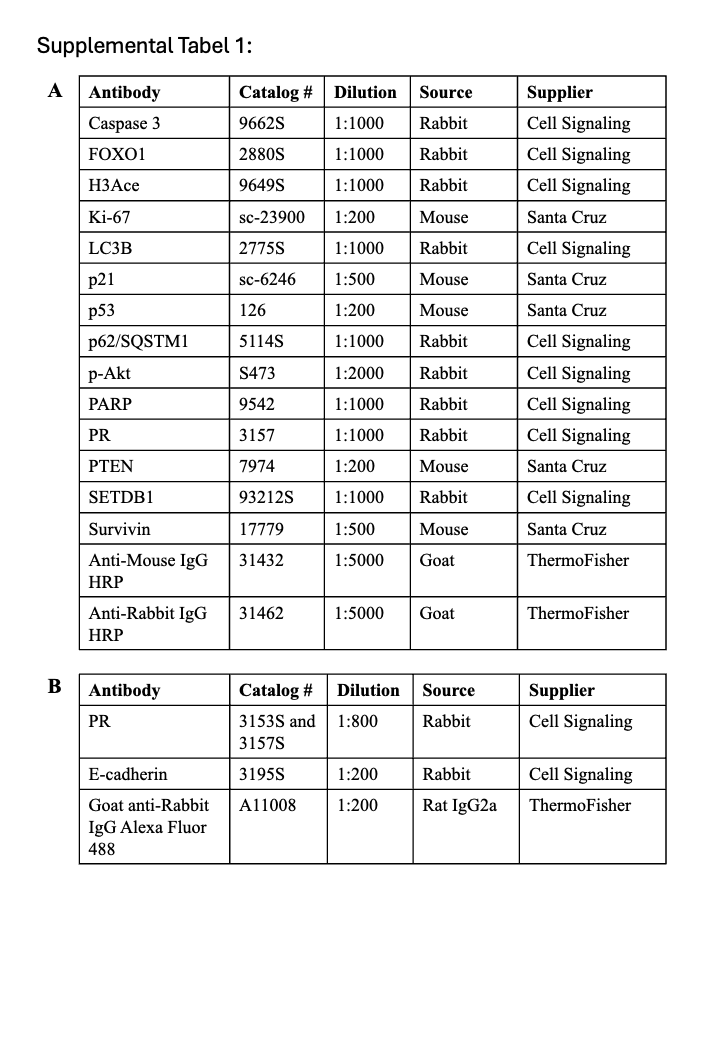
**Supplemental Tabel 1.** Antibodies used in Western Blot analysis and immunofluorescence.

(A) Antibodies used in Western Blot. B) Antibodies used in immunofluorescence labeling of spheroids.
